# Screening Immunotherapy Targets to Counter Radiation-Induced Neuroinflammation

**DOI:** 10.1101/2022.08.23.505001

**Authors:** Sadhana Sharma, Christina Fallgreen, Michael M. Weil, Anushree Chatterjee, Prashant Nagpal

## Abstract

Galactic cosmic rays (GCR) in space induce increase in cerebral amyloid-β levels and elevated levels of microgliosis and astrocytosis, causing accelerated neurodegeneration from this increased neuroinflammation. Even exposure to low-levels of high-Z high-energy (HZE) radiation (50 cGy) has been shown to induce biochemical and immunohistochemical changes in short-term leading to degradation in cognition, motor skills, and development of space-induced neuropathy. There is lack of effective neuroinflammation countermeasures, and current experimental therapies require invasive intracerebral and intrathecal delivery due to difficulty associated with therapeutic crossover between blood-brain barrier. Here, we present a new countermeasure development approach for neurotherapeutics using high-throughput drug-discovery, target validation, and lead molecule identification with nucleic acid-based molecules. These Nanoligomer™ molecules are rationally designed using a bioinformatics and AI-based ranking method and synthesized as a single-modality combining 6-different design elements to up- or down-regulate gene expression of target gene at will, resulting in elevated or diminished protein expression of intended target. This platform approach was used to perturb and identify most effective upstream regulators and canonical pathways for therapeutic intervention to reverse radiation-induced neuroinflammation. The lead Nanoligomer™ and corresponding target granulocyte-macrophage colony-stimulating factor (GM-CSF) were identified using *in vitro* cell-based screening in human astrocytes and donor derived peripheral blood mononuclear cells (PBMCs) and further validated *in vivo* using a mouse model of radiation-induced neuroinflammation. GM-CSF transcriptional downregulator Nanoligomer 30D.443_CSF2 downregulated proinflammatory cytokine GM-CSF (or CSF2) using simple intraperitoneal injection of low-dose (3mg/kg) and completely reversed expression of CSF2 in cortex tissue, as well as other neuroinflammation markers. These results point to the broader applicability of this approach towards space countermeasure development, and potential for further investigation of lead neurotherapeutic molecule as a reversible gene therapy.

## INTRODUCTION

Space explorations are on the rise, not only for scientific explorations, but more recently for space tourism. As space missions get longer and progress away from the lower Earth orbit and thus from Earth’s protecting magnetic shielding, the radiation exposures that astronauts and space tourists will need to face will expand to the full galactic cosmic ray spectrum and solar particle events.^1^ Exposure to even low-levels of high-Z high-energy (HZE) GCR radiation (50 cGy) has been shown to induce increase in cerebral amyloid-β (Aβ) levels.^2,3^ Aβ acts as a pro-inflammatory cytokine^4^ inducing biochemical and immunohistochemical changes leading to increased levels of microgliosis and astrocytosis, protein misfolding, and neurodegeneration.^5–11^ Hypothesized reasons for the accelerated neurodegeneration are centered around a combination of oxidative stress and mitochondrial dysfunction leading to increased inflammation,^12^ oxidative species induced protein misfolding and plaque buildup,^2,12^ microgravity induced neuropathy,^3^ radiation induced damage to DNA-repair mechanisms, and simultaneous development of other pathologies all leading to accelerated neurodegeneration.^13–17^ While the specific mechanisms of neurodegeneration and space-induced neuropathology are being studied in simulated conditions on earth for space medicine, protection of astronauts on spaceflights, clearly documented simultaneous degradation in cognition and motor skills,^18^ and development of space-induced neuropathy outlines the urgent need for rapid countermeasure development.

There are currently no approved radiation-induced neurodegeneration countermeasures available for the treatment. Specifically, cerebrovascular sub-syndrome is considered incurable. Recombinant granulocyte-macrophage colony-stimulating factor (rhGM-CSF, sargramostim, Leukine) and human granulocyte colony-stimulating factor (rhG-CSF, filgrastim, Neupogen) are the only two FDA-approved countermeasures for accelerating hematologic recovery.^19,20^ However, GM-CSF’s proinflammatory role has been implicated in several autoimmune diseases, including rheumatoid arthritis and multiple sclerosis.^21,22^ Its potential role in inducing blood-brain barrier (BBB) permeability ^21,23–25^ and neuroinflammation,^22^ leads to astrocytosis and microgliosis and neurodegenerative diseases make GM-CSF a good countermeasure/downregulation target. ^21,26,27^ rather than an upregulated recombinant protein, to counter radiation-induced neuropathy and neuroinflammation. Recombinant protein-based countermeasures like Sargramostim and Filgrastim, suffer from limitations associated with potential protein degradation and denaturation, especially during space travel. Despite remarkable progress in the field of nucleic acid therapeutics based on microRNAs, small interfering RNAs, long non-coding RNAs, and deactivated CRISPR-Cas9 technology in the recent years, several challenges associated with their delivery, stability, internalization, and target specificity need to be addressed for their successful clinical translation and development as radiation countermeasures.^28,29^ Moreover, technologies like deactivated CRISPR-Cas9 require tedious cloning and optimization that are time-consuming and require specific skills that may not be available on longer space flight missions. Hence, there is an urgent need to develop new therapeutic modalities that can address above-mentioned challenges, and quickly develop, fine tune, and deliver precision medicine during or after space travel.

Sachi has developed Nanoligomer™ technology, a high-precision tool that generates sequence-specific, nano-biohybrid molecules called Nanoligomers™ to address this unmet need.^30^ A Nanoligomer has six design elements (proprietary) and can be designed to either up- or down-regulate any desired gene by either binding to its mRNA or DNA. The nucleic acid-binding domain of Nanoligomers is peptide nucleic acid (PNA), which is a synthetic DNA-analog where the phosphodiester bond is replaced with 2-N-aminoethylglycine units.^31^ Nanoligomers offer improved stability, facile delivery and internalization, and superior target specificity compared to existing nucleic acid therapeutics.^32^ In addition, they modulate related gene network targets ultimately resulting in pathway modulation. Previously, we have demonstrated that Nanoligomer technology is very effective in identifying lead targets and molecules for reversing radiation-induced immune dysfunction.^30^

Based on the literature,^18,33–35^ we identified six key targets, viz., Granulocyte-Macrophage Colony-Stimulating Factor (GM-CSF), Interleukin-6 (IL-6), Interleukin-1β (IL-1β), Tumor Necrosis Factor alpha (TNF-α), Tumor Necrosis Factor receptor 1 (TNFR1), Interleukin10 (IL-10) and screened them for developing best-in-class countermeasures for radiation-induced neuroinflammation. Overexpression of pro-inflammatory mediators such as GM-CSF, IL-6, IL-1β, or TNF-α in the radiation-exposed brain indicate a greater proportion of activated microglia, and hence likely a greater degree of neuroinflammation.^36,37^ While GM-CSF (Sargramostim, Leukine) delivery is currently the first line of treatment for hematologic recovery after accidental radiation exposure,^19,20^ its proinflammatory role has been implicated in several autoimmune diseases, including neurodegenerative diseases like multiple sclerosis.^21,22^ Due to multiple potential pathways such as increasing BBB permeability^21,23–25^ and neuroinflammation,^22^ increased GM-CSF levels have been linked to astrocytosis and microgliosis and neurodegenerative diseases.^21,26,27^ IL-6 plays a critical role in neuroinflammation and blocking IL-6 signaling with a monoclonal antibody against its receptor (Tocilizumab) has been approved for the treatment of several inflammatory diseases.^36–38^ IL-1β is another key mediator that drives neuroinflammation,^34,39^ and its inhibitor in the form of recombinant protein is approved for treating rheumatoid arthritis and is in clinical trials for treating other chronic inflammatory diseases.^40,41^ TNFα is an important therapeutic target for developing therapies for controlling inflammation.^42^ Several recombinant proteins and antibodies that bind to and neutralize TNF have been investigated for various inflammatory diseases. TNF-α activates two types of receptors-TNFR1 and TNFR2. TNFR1 promotes apoptosis and activates NF-κB, which leads to proinflammatory cytokine production and glial activation, resulting in neuroinflammation and neuronal death.^4,43^ Together, IL-1 β and TNF-α have been shown to cause neuronal death synergistically by the direct effects of these cytokines on neurons or indirectly by glial production of neurotoxic substances.^44^ In addition, they induce death of astrocytes that results in the transient production IL-6 and other inflammatory mediators.^45^ Interleukin 10 (IL-10) is an anti-inflammatory cytokine produced by multiple immune cell types including regulatory T cells (Tregs) and plays a prominent role in limiting immune-mediated inflammation during radiation exposure, infection, allergy, or autoimmunity.^46^

To screen and rank these targets and identify the best-in-class and potential first-in-class therapeutic molecule, we applied Sachi’s Nanoligomer platform to regulate the above mentioned neuroinflammation relevant targets to identify the most effective upstream regulator for developing countermeasure for radiation-induced neuroinflammation.

## RESULTS AND DISCUSSION

Sachi’s Nanoligomer technology enables both up- and downregulation of any gene of interest by following the four steps-Design, Build, Test and Learn-of synthetic biology (Fig. 1A). Nanoligomer™ is a nano-biohybrid molecule incorporating 6-different design elements (proprietary).^30^ The nucleic acid-binding domain of Nanoligomers is peptide nucleic acid (PNA), which is a synthetic DNA-analog where the phosphodiester bond is replaced with 2-N-aminoethylglycine units.^31^ Peptide Nucleic Acids (PNAs) demonstrate stronger hybridization and target specificity compared to binding of naturally occurring RNA or DNA^47^ and can withstand both nucleases and proteases, leading to increased stability in human blood serum and mammalian cellular extracts.^48^ Nanoligomer downregulators act either *via* targeting of mRNA to inhibit RNA stability and consequently protein expression or *via* targeting of DNA.^30,32^ Nanoligomer upregulators trigger transcriptional activation by binding at the respective promoter region and attachment of specific domains that recruit transcriptional activators (Fig. 1B).

**Figure 1.**
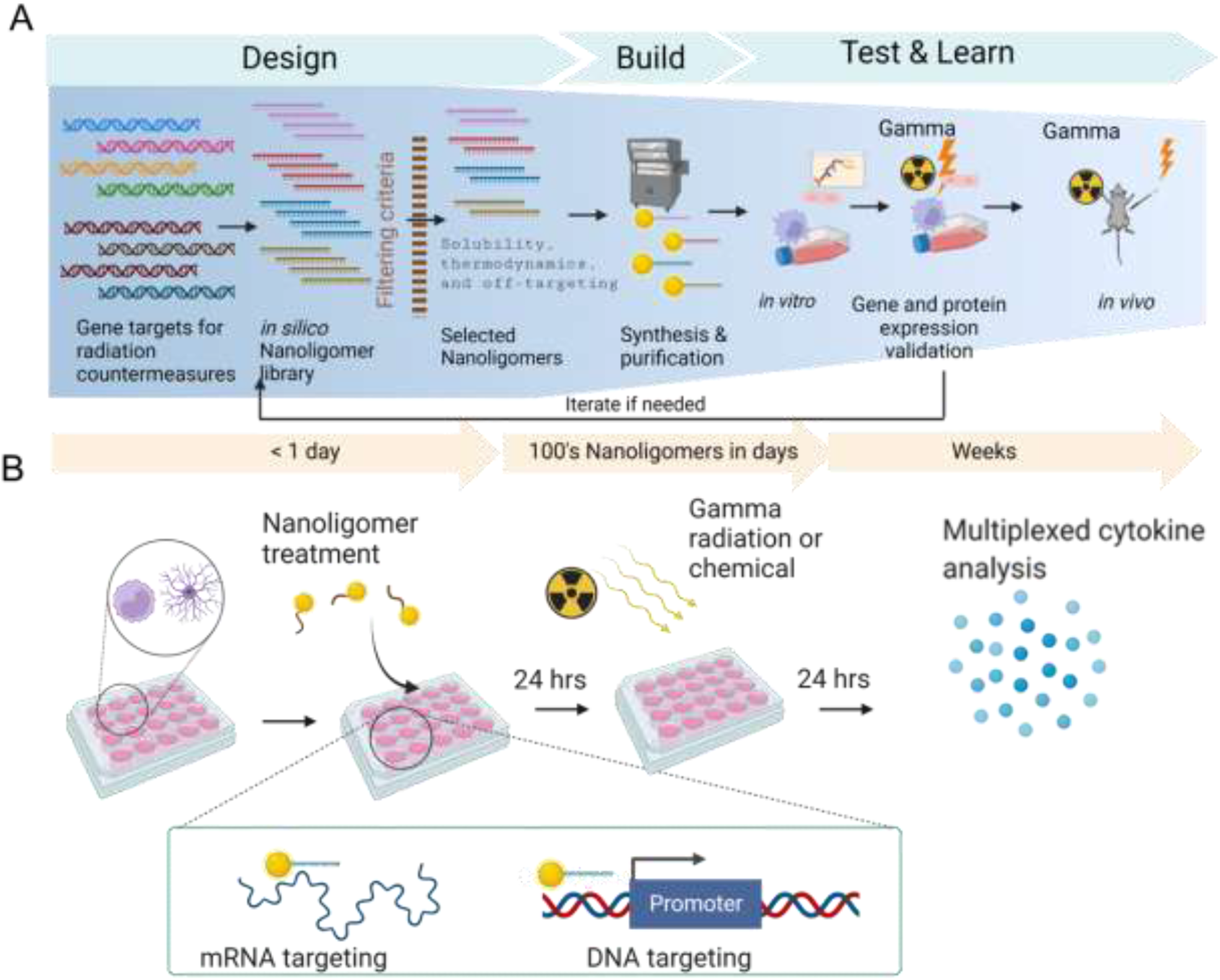
Design and evaluation of Nanoligomer based countermeasures for radiation-induced neuroinflammation. (A) Process flow for Nanoligomer platform follows steps of Design, Build, Test and Learn. An in silico Nanoligomer library is created for each selected gene target and all candidates are evaluated and selected through a four-step process. First, gene target regions (RNA and/or DNA) are identified based on factors such as sequence composition and predicted transcription or translational regulatory regions. These seed regions are then evaluated for expected synthesis and solubility considerations followed by filtering based on undesirable features such as self-complementary and direct match or multiple mismatch off-targets. The fourth step is a thermodynamic analysis using a Naïve Bayes classifier for target vs off-target binding and optimization of possible on-target effectiveness. Best Nanoligomer designs are synthesized, purified, and screened in astrocytes and irradiated donor derived PBMCs via multiplex cytokine panel. Finally, an identified Lead Nanoligomer candidate is tested in a murine model of radiation-induced neuroinflammation and evaluated by cytokine panel for its ability to reduce neuroinflammation. (B) Experimental in vitro set up to evaluate top candidate Nanoligomers based countermeasures for neuroinflammation. Inset shows the two mechanisms of action by which Nanoligomers can regulate gene expression. Left: Nanoligomers target mRNA to regulate RNA stability or translation. Right: Nanoligomers target genomic DNA to regulate transcription.

### Process flow for Sachi’s Nanoligomer Platform

To design Nanoligomers, the sequences of the identified gene targets are used as inputs to Sachi’s bioinformatics toolbox (Fig. 1A). An *in silico* Nanoligomer library is created for each selected gene target and all candidates are evaluated and selected through a four-step process. First, gene target regions (RNA and/or DNA) are identified based on factors such as sequence composition and predicted transcription or translational regulatory regions. Second, these seed regions are then evaluated for expected synthesis and solubility considerations to maximize therapeutic efficacy. Next, filtering based on undesirable features such as self-complementary and direct match or multiple mismatch off-targets (up to two base pair mismatches) is done for each candidate along the human genome. The fourth step is a thermodynamic analysis using a machine-learning approach based on Naïve-Bayes classifiers^49–52^ for target vs off-target binding and optimization of possible on-target effectiveness. For each target, this process is iterated several times to design Nanoligomers targeting different sites on the genome. Scoring and ranking of the bioinformatics generated *in silico* Nanoligomer library was achieved as described in detail previously.^30^ Briefly, we ranked the immunotherapy targets and respective Nanoligomer molecules using a score based on potential off-targets, binding specificity, etc. The first weighing coefficient for target specificity (*ω*_*iMM*_) was the ratio of measured dissociation constant (*K*_*D*_) with respective mismatches 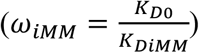. The score was assigned using product of the mismatch throughout human genome (*C*_*iMM*_) and the respective weighing coefficient. For binding affinity scores (*ω*_*TM*_), we used respective melting temperatures (*T*_*M*_) to obtain coefficient as 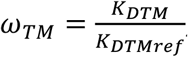, where dissociation constant is calculated from melting temperatures using 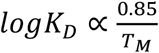 and as 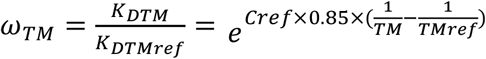. Combining the effects of binding specificity and binding affinity provides an overall ranking 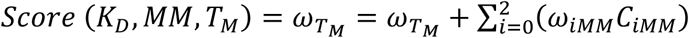.

After completing the design process, top Nanoligomer designs (at least three per target-one per genomic location) incorporating all 6-different design elements were synthesized as a single-modality using high throughput peptide synthesizer and purified (Fig. 1A). In addition, a Missense Nanoligomer (designed with no homology to any gene in the specific genome at DNA/ mRNA level) was designed to rule out any possible non-specific effects. Next, Nanoligomers were screened *in vitro* using appropriate cell models (astrocytes and irradiated donor derived PBMCs in this study, see details in next section) to rank and validate by multiplex cytokine panel (Fig. 1A, B). Finally, *in vitro* validated Lead Nanoligomer candidate was tested in a murine model of radiation-induced neuroinflammation and evaluated by multiplex cytokine assay for its ability to mitigate neuroinflammation (Fig. 1A).

### Assessing Relative Importance/Efficacy of Target in Neuroinflammation

The relative importance and efficacy of the selected targets, viz., GM-CSF, IL-6, IL-1 β, TNF-α, TNFR1, and IL-10, in neuroinflammation was assessed based on *in vitro* screening using two neuroinflammation linked cell models, namely, astrocytes and irradiated donor-derived PBMCs, followed by multiplexed 65-cytokine analysis (Fig. 1A, B; Fig. 2). Cytokine levels were normalized with respect to no treatment and significance was calculated using a t-test between replicates. Any cytokine that showed broad expression changes across multiple treatments were removed from analysis to elucidate gene specific effects. Choice of the cell model-target combination was guided by the basal (gene/protein) expression level in the cells.

**Figure 2.**
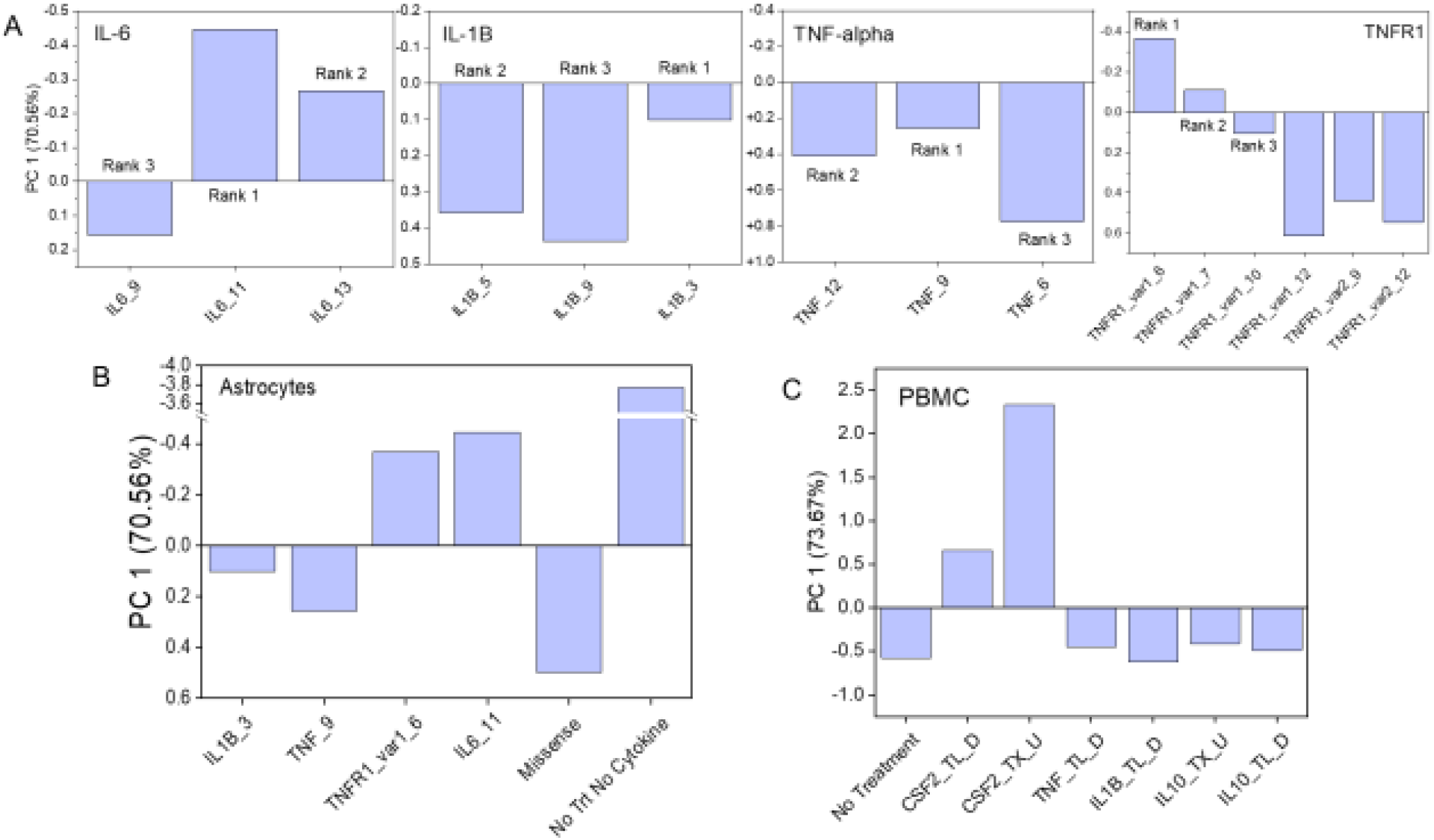
Principal component analysis of human astrocytes and irradiated PBMC cytokine expression data for various Nanoligomer treatments. (A) PC1 data for multiple Nanoligomer designs targeting IL-6, IL-1B, TNF-alpha, and TNFR in human astrocyte screens. (B) PC1 data for best Nanoligomer designs identified based on astrocyte screen for IL-6, IL-1B, TNF-alpha, and TNFR1 targets along with missense and no Nanoligomer/ no cytokine treatment controls. (C) PC1 data for up (transcriptional) and down (translational) regulator Nanoligomer targeting CSF2, TNF-alpha, IL-1B, and IL-10 based on PBMC screens at 3 Gy radiation and no Nanoligomer treatment control. U: upregulator, D: downregulator; Var1: variant 1, Var2: variant 2 TL: translational; TX: transcriptional. PCA is based cell culture and cytokine data obtained from three biological replicates.

Astrocytes play a key role in brain homeostasis, maintaining the blood-brain barrier (BBB), and regulating inflammatory responses in the central nervous system.^53^ In the presence of appropriate stimuli, they can be either neurotoxic (A1-phenotype) or neuroprotective (A2-phenotype). An alteration in the relative population of these phenotypes (more A1-phenotype, less A2-phenotype) has been linked to neuroinflammation and neurodegenerative diseases.^53^ PBMCs, the major immune cells in the human body whose infiltration into the brain has been known to drive neuroinflammation^54^, have been used extensively as an *in vitro* test platform for immune modulation and inflammation with and without radiation exposure.^30,55,56^ At least three Nanoligomers per target were designed, synthesized as a single-modality (incorporating all 6-different design elements) using a high throughput peptide synthesizer, and purified.

Human primary astrocytes were used for screening four targets, viz., IL-6, IL-1 β, TNF-α, and TNFR1, using multiple translational (targeting mRNA) Nanoligomer downregulators (Fig. 2A,B). For TNFR1, Nanoligomers targeting variant 1 and 2 (listed in the NCBI database) were used. Briefly, astrocytes were first pre-treated with the gene-specific Nanoligomers (with or without cytokine cocktail) for 24 hours, and following a further 24 hours, media supernatants were collected and analyzed for secreted protein/cytokine expression (see methods for details). Cytokine cocktail (TNF-α, IL-1α and C1q) induces astrocytes towards neuroinflammatory A1-phenotype.^4,57^ The high-dimensional (65-plex) cytokine data was normalized to their respective cytokine levels in non-treated astrocytes and transformed using principal component analysis (PCA). PCA is unbiased technique, like clustering, that can be used to reduce dimensionality for a complex data set to find trends and patterns.^58^ Scree plot (Fig. S1A) revealed that most of the variance (70.56%) in the data is captured by principal component 1 (PC1). Based on the PC1 scores (more negative values imply better efficacy of Nanoligomer downregulator), we ranked the various Nanoligomer designs for each gene target (Fig. 2A). Next, we selected the top-ranked (Rank 1) Nanoligomer inhibitor for all the targets and compared their PC1 with missense and “No treatment no cytokine” control (Fig. 2 B). Based on this analysis, we made the following assessments: 1) IL-1B_3, TNF_9, TNFR1_var1_6, and IL6_11 were the best Nanoligomers for their respective targets. 2) The Target Ranking was: IL-6 (Rank 1), TNFR1 (Rank 2), TNF-α (Rank 3), and IL-1 β (Rank 4). The PCA and ranking analysis suggested IL-6 was the most promising gene target for neuroinflammation, since it was the farthest from the missense control that represents diseased state, and no treatment with cytokine cocktail-induced neuroinflammation) and closest to the no treatment no cytokine control (that represents normal healthy state, no neuroinflammation). However, IL-6 is a pleiotropic cytokine that fulfils multiple contrasting functions: 1) essential homeostatic functions, which include immune cell proliferation and differentiation as well as metabolic functions, 2) pro-inflammatory actions due to dysregulated activity. Therefore, sparing the homeostatic functions of IL-6 in order to avoid serious long term side effects is crucial when targeting IL-6.^59^ Although monoclonal antibodies against IL-6 or IL-6 receptor (IL-6R) and Janus kinases (JAK) inhibitors developed for the treatment of autoimmune diseases such as rheumatoid arthritis have shown promise in clinical trials, their use resulted in increased risk of bacterial and viral infections, essentially compromising the immune defense of the brain and exposing the host to a range of debilitating infections and diseases. Therefore, the broad role of IL-6 in mediating several key physiological pathways in literature, made it unsuitable for broadband downregulation to treat neuroinflammation and radiation-induced neuropathy.^59^

For further screening of targets for radiation-induced inflammation, we used irradiated PBMCs to screen four targets, viz., GM-CSF, IL-1 β, TNF-α, and IL-10 (Fig. 2C). Top ranking Nanoligomers for IL-1 β and TNF-α based on astrocyte screening were also used in PBMC-based screening. For GM-CSF and IL-10, multiple Nanoligomers-translational (targeting mRNA) downregulators and transcriptional (targeting DNA) upregulators-were examined. Briefly, PBMCs pre-treated (24 hours) with the gene-specific Nanoligomers were exposed to gamma-radiation at pre-determined dose (3 Gy), and after another 24 hours media supernatants were collected and analyzed for secreted protein/cytokine expression. Based on Scree plot (Fig. S2B), most of the variance (73.67%) in the data was captured by principal component 1 (PC1). So, Nanoligomers and targets were ranked based on PC1 scores (higher positive values imply better efficacy of Nanoligomer upregulator, more negative values imply better efficacy of Nanoligomer downregulator). Fig. 2C shows PC1 scores for Nanoligomers (up/down, translational/transcriptional regulators) for all the targets as compared to irradiated (3 Gy), no treatment control PBMCs (representing radiation-exposed neuroinflammatory state). Based on this analysis, we made the following assessments: 1) Target ranking: CSF2 (GM-CSF, Rank 1), IL-1 β (Rank 2), TNF-α ∼ IL-10 (Rank 3), suggested CSF2 (GM-CSF) as the clear lead gene target for neuroinflammation as PC1 values for all CSF2 Nanoligomer designs are the farthest from the irradiated (3 Gy), no treatment control as compared to all other targets. 2) Comparison of CSF2 (GM-CSF) targeting translational (targeting RNA) downregulators and transcriptional (targeting DNA) upregulators under various radiation exposure conditions suggested that transcriptional regulators are more effective in downregulating the intended GM-CSF target.

Based on PCA analysis of multiplexed 65-plex cytokine data, we decided to move forward with CSF2 (GM-CSF) as the lead target. GM-CSF initiates upstream inflammatory gene expression in monocytes and their progeny (multilineage immune upregulation has been observed, ^60–63^), which further effects GM-CSF-dependent pathogenesis in the inflamed tissue. GM-CSF or GM-CSFR blockade with monoclonal antibodies has also been investigated for several inflammatory diseases, such as rheumatoid arthritis, asthma and COVID-19, and has shown encouraging results in early phase clinical trials. ^64^ This also suggests that the downregulation of GM-CSF could be a useful strategy for reducing or reversing radiation-induced neuroinflammation.

### Nanoligomer GM-CSF regulators modulate GM-SCF expression and system level cytokine response

To validate CSF2 (GM-CSF) as the lead target for neuroinflammation, we further analyzed GM-CSF protein expression data for the lead GM-SCF upregulator CSF2-U (CSF2_TX_U, Nanoligomer library name: SB201.1U.1 CSF2) and downregulator CSF2-D (CSF2_TL_D Nanoligomer library name: SB1D.431 CSF2). It is worth mentioning here that GM-CSF upregulator is transcriptional i.e. it targets host DNA and GM-CSF downregulator is translational i.e. it targets mRNA, to further parse out the efficacy of transcriptional vs translational downregulation. These Nanoligomers were tested with (3 Gy) or without radiation exposure (0 Gy) using irradiated PBMCs. We observed that CSF2-U significantly upregulated GM-SCF protein expression w.r.t. no treatment control (for 3 Gy exposure) (Fig. 3A). Also, CSF2-D significantly downregulated of GM-CSF protein expression with respect to (w.r.t.) missense control (for no radiation exposure) (Fig. 3B). Previously, CSF2-U and CSF2-D Nanoligomers were validated for their up- and down regulatory potential at mRNA level with HepG2 (human hepatocellular carcinoma, ATCC) cells using real-time quantitative polymerase chain reaction (RT-qPCR). As in Fig. S2, *csf2*gene was significantly upregulated by CSF2-U.

**Figure 3.**
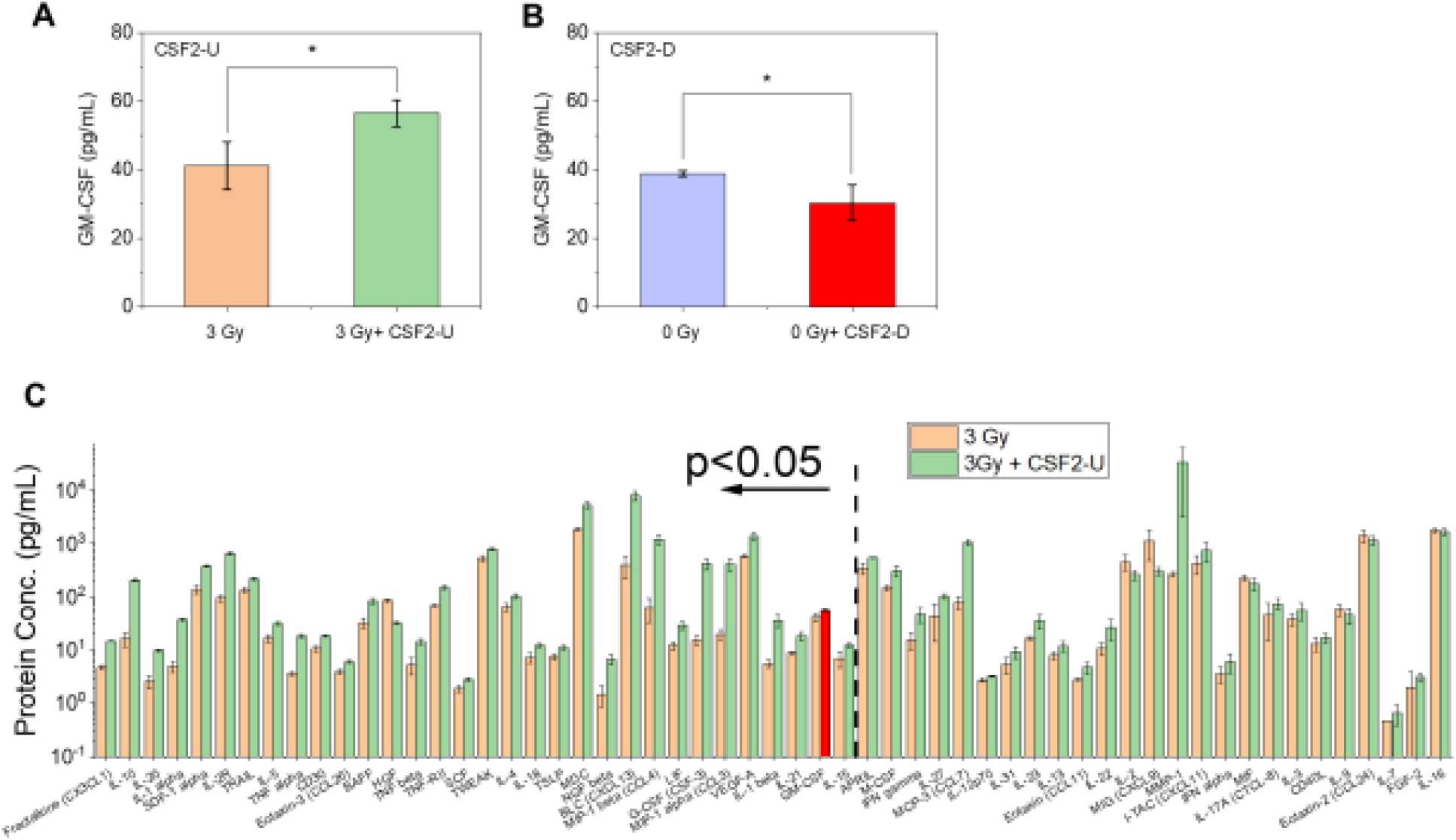
GM-CSF regulators modulate cytokine/ protein expression in PBMCs (A) GM-CSF upregulator (CSF2-U) increases its expression, 3 Gy radiation exposure (B) GM-CSF downregulator (CSF2-D) decreases its expression, no radiation exposure (C) System level cytokine/protein response to GM-CSF upregulator (CSF2-U). Data shown is an average of three biological replicates, and error bars represent standard deviation.

Fig. 3C shows the system level cytokine response of GM-CSF upregulator (CSF2-U) after 3 Gy gamma radiation exposure (data normalized to 3 Gy, no treatment vehicle control). Out of 65 secreted proteins, 32 were differentially expressed (p<0.05), out of which, 31 proteins (GM-CSF, G-CSF(CSF3), TNFα, TNFβ, IL-4, IL-5, IL-1A, IL1b, IL-10, IL-15, IL-18, IL-20, IL-21, LIF, TSLP, Fractalkine(CX3CL1), Eotaxin-3, SDF-1α, BLC (CXCL13), MIP-1α (CCL3), MIP-1β (CCL4), MDC (CCL22), SCF, VEGF-A, NGFβ, TNF-RII, TWEAK, TRAIL, CD30, BAFF, IL-2R (CD25)) were upregulated, and the remaining one-growth factor HGF2-was downregulated. CSF2-U upregulated GM-CSF expression as expected, but it also upregulated closely associated G-CSF, which also plays important role in hematologic recovery. ^65^ Interleukin-3 (IL-3), which clusters with GM-CSF and G-CSF (STRING database) and plays important role in hematopoiesis as well as in amplifying acute inflammation (a potential therapeutic target in sepsis), was also upregulated but not significantly. Growth factors, viz., SCF and NGFβ were upregulated. Stem cell factor (SCF) regulates embryonic and adult hematopoiesis and thus favors hematologic recovery after radiation exposure^66^. NGFβ, secreted by PBMCs, is known to delay neurodegeneration in the central nervous system.^67^

We observed an upregulation of both pro- and anti-inflammatory cytokines in PBMCs, suggesting that the balance between the two would guide the actual inflammatory response. Pro-inflammatory response was primarily mediated by cytokines of IL-1 family (IL-1α, IL1β), TNF family (TNF α, TNFβ), and chemokines (MIP-1α (CCL3), MIP-1β (CCL4)), and supported by GM-CSF, G-CSF(CSF3), IL-15, IL-18, IL-20, IL-21, TSLP, TWEAK, TRAIL, CD30 and BAFF upregulation.^68^ IFNγ, a proinflammatory cytokine known for its role in inflammation and autoimmune disease, was not significantly upregulated. Although we did not observe any changes in IL-6, which plays a critical role in neuroinflammation, we did observe upregulation of LIF, a key member of the IL-6 family of cytokines. Upregulation of VEGFα and downregulation of HGF2 indicate pro-inflammation. Several chemokines, namely, Fractalkine (CX3CL1), Eotaxin-3 (CCL26), SDF-1α (CXCL12), BLC (CXCL13) and MDC (CCL22) were upregulated suggesting the activation and migration of immune cells. Anti-inflammatory response was primarily mediated by IL-10, IL-4, IL-5, and TNF-RII upregulation. IL-10 is an anti-inflammatory cytokine produced by multiple immune cell types including regulatory T cells (Tregs) and plays a prominent role in limiting immune-mediated inflammation during radiation exposure, infection, allergy, or autoimmunity.^46^ IL-10, IL-4, and IL-5 have potent anti-inflammatory properties and can effectively suppress the production and function of pro-inflammatory cytokines and their receptors at multiple levels.^68^

The soluble Tumor necrosis factor receptor 2 (TNF-RII), which binds to tumor necrosis factor-alpha (TNFα), reduces pro-inflammatory effects of TNFα, activates and expands Tregs, and favors neuroprotection.^69,70^ Interleukin-2 receptor subunit alpha (IL-2Rα or CD25) was also upregulated. It plays a key role in regulating immune system and tolerance *via* regulation of Forkhead box P3 (FoxP3), a transcriptional factor, which is crucial for the development and inhibitory function of regulatory T cells (Tregs).^71^ Co-expression of TNF-RII and CD25 has been associated with increased number of the functional CD4^+^FOXP3^+^ Tregs in human peripheral blood ^72^ Overall, the cytokine data for GM-CSF upregulator (CSF2-U) suggests hematologic recovery but does not give a clear indication of overall pro- or anti-inflammatory response. In addition, *in vitro* screening for all the targets indicated that the transcriptional regulators are more efficient in regulating the target in the desired direction as compared to the translational regulators. Therefore, we decided to design a transcriptional downregulator for GM-CSF for *in vivo* testing.

### A low-dose of GM-CSF transcriptional downregulator administered intraperitoneally reverses radiation-induced neuroinflammation in a mouse model

To evaluate transcriptional GM-CSF downregulating Nanoligomer (as identified using the Nanoligomer platform) in an *in vivo* mouse model of radiation-induced neuroinflammation (Fig. 4A), we designed and synthesized Nanoligomer 30D.443_CSF2 targeting mouse GM-CSF/ CSF2 gene using Sachi’s Nanoligomer pipeline. Next, we evaluated GM-CSF downregulator (30D.443_CSF2) in *vivo* using eight-week-old female C57BL/6NCrl mice (Charles River Laboratories). First, we assessed the immunogenicity and toxicity of 30D.443_CSF2 Nanoligomer by examining the serum cytokines known to be involved in several pathological immune responses. The responses considered were acute-phase response (IL-1β, IL-6, TNFα), Cytokine storm/release (IL-2, IL-6, IL-10, IFNγ, TNFα), Hemophagocytic syndrome (IFNγ, IL-1β, IL-6, TNFα), Neutrophilic inflammation (MIP-1α, TNFα), Systemic inflammatory response syndrome (IL-6, CCL2, TNF-α), and Th1 (IFNγ, IL-2, IL-12) and Th2 (IL-4, IL-5, IL-6, IL-10, IL-13) response.^73^ For all cytokines indicated in these responses, we observed no significant changes in their serum levels with Nanoligomer treatment compared to controls (Fig. S5). Most remained below the limit of quantification in our assays which ranges from 0.09 pg/mL for IL-3 to 2.4 pg/mL for CCL2 (data points below LOQ are represented at 0.5LOQ). This lack of blood cytokine biomarkers of inflammation and *in vivo* toxicity supports the safety of Nanoligomers and the lack of broad, untargeted immune modulation.

**Figure 4.**
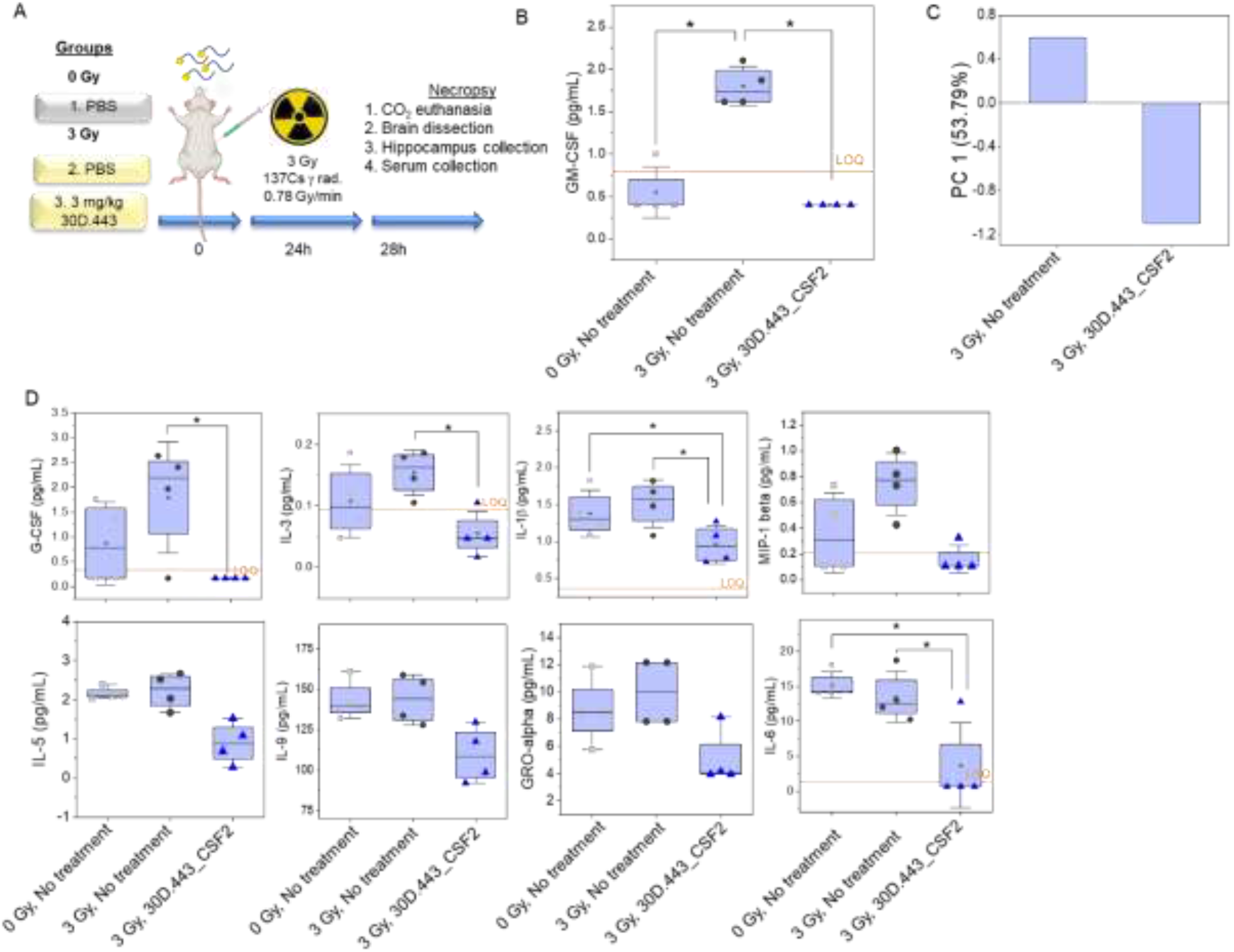
*In vivo* evaluation of GM-CSF downregulator (30D.443_CSF2) in a mouse model of radiation. A) Schematic of the in vivo mice radiation model used to assess the efficacy of the identified best-in-class neurotherapeutic30D.443_CSF2. B) 30D.443_CSF2 downregulates GM-CSF protein expression in mouse cortex (p<0.01) after 24 hours of Nanoligomer treatment and 3 Gy radiation exposure. C) Principal component analysis shows a distinction between irradiated 30D.443_CSF2 treated and untreated irradiated mice based on PC1 that accounts for 53.79% of data variability. D) Cytokine expressions for G-CSF, IL-3, IL-1β, MIP-1β (CCL4), IL-5, IL-9, GRO-alpha, and IL-6 (p<0.05) upon treatment with 30D.443_CSF2 with (3 Gy) or without (0 Gy) radiation exposure. Orange line represented the limit of quantification (LOQ) of the assay, symbols represent individual animals (n=4 mice), horizontal bar is the average and error bars are +/-one standard deviation.

Next, we evaluated GM-CSF downregulator (30D.443_CSF2) in *vivo* using mouse model of radiation. The mice were split into three treatment groups (4 mice/ group). These groups were administered either vehicle negative control (Group 1 and 2) or 30D.443_CSF2 Nanoligomer (Group 3) via intraperitoneal (IP) injection. Our previous *in vivo* mouse studies have shown that Nanoligomer provides a facile route for delivery through peritoneum. The Nanoligomer molecules not only cross the BBB, but also have low K_D_ (dissociation constant), facilitating excellent therapeutic action even at low doses *in vivo*.^32^ IP injection of GM-CSF downregulator showed Nanoligomer in the brain within 1 hour of injection (data not shown). Twenty-four hours post injection (hpi), Group 2 (vehicle) and group 3 (30D.443_CSF2) were exposed to a total dose of 3 Gy gamma total body irradiation (Fig. 4A). Due to the Nanoligomer treatment, mice did not show any signs of distress. At twenty-eight hours post injection (hpi), mice were euthanized by carbon dioxide, blood was immediately collected by direct cardiac draw and processed to serum, brains were dissected, and cortex tissues were flash frozen.

Flash frozen mouse cortex tissues were processed and analyzed for cytokines and chemokines using 36-Plex Mouse ProcartaPlex Panel 1A (ThermoFisher Scientific, see methods for details), which also includes the targeted CSF2/GM-CSF. The basal expression of CSF2/GM-CSF in the mouse cortex was below the level of detection (Fig. 4B). However, CSF2 was significantly (*p*<0.01) elevated (∼3-fold) as a result of 3 Gy radiation exposure (3 Gy, vehicle) compared to no radiation (0 Gy, vehicle). Treatment with 30D.443_CSF2 Nanoligomer in the irradiated state (3 Gy, 30D.443_CSF2) brought the CSF2 expression back to basal level (*p*<0.01) (Fig. 4B), demonstrating that Sachi’s GM-CSF downregulator (30D.443_CSF2) effectively reduces CSF2/GM-CSF protein in the mouse cortex.

Principal component analysis of the cytokine data revealed that most of the variance (53.79%) in the data is captured by principal component 1 (PC1) (Scree plot, Fig. S6). PC2, PC3, and PC4 capture only 10.59%, 9.79%, and 8.69% of variability, respectively. PC1 plot (Fig. 4C) clearly shows a distinction between 30D.443_CSF2 treated and untreated radiation-exposed mice, suggesting that Nanoligomer treatment results in broad alterations in the cytokine profile in mouse cortex.

Additionally, we looked at the twenty closest functional and physical related protein associates to GM-CSF (CSF2) in mouse (STRING database)^74^ of which seven were present in the 36-plex panel: G-CSF (CSF3), IL-3, IL-4, IL-5, IL-6, IL-15, and TNFα. Treatment with 30D.443_CSF2 Nanoligomer in the irradiated state (3 Gy, 30D.443_CSF2) significantly downregulated the expression of G-SCF, IL-3, IL-1β, IL-5, IL-9, GROα (CXCL1), and MIP-1β (CCL4), either back to or lower than the basal level (p<0.01) (Fig. 4D), demonstrating the capability of Sachi’s GM-CSF downregulator (30D.443_CSF2) in controlling radiation-induced neuroinflammatory milieu. In addition, IL-6, TNFα, and CCL2 were significantly lower in the cortex with 3 Gy, 30D.443_CSF2 treatment than in no radiation no treatment condition (0 Gy, vehicle) (Fig. S7). These results establish that GM-CSF downregulator (30D.443_CSF2) is effective in reducing radiation-induced neuroinflammation in the selected mouse model of radiation. Cytokine levels in the mouse cortex for the remaining cytokines (ENA-78 (CXCL5), Eotaxin (CCL11), IP-10 (CXCL10), MIP-1α (CCL3),, IL-10, MIP-2α (CXCL2), RANTES (CCL5), IFNγ, IL-1α, IL-4, IL-15/IL-15R, IL-17A (CTLA-8), IL-18, IL-28, IL-31, LIF, MCP-3 (CCL7), M-CSF) with and without 3 Gy radiation and treatment with 30D.443_CSF2 are shown in Fig. S7.

Taken together, the safety profile and network wide immune effect of 30D.443_CSF2 supports further investigation of this Nanoligomer as well as the platform in general for neuroinflammation countermeasure development. Further studies for neuroinflammation should explore variable dosing regimens, alternate route of administration such as intranasal, as well as multiplexing with other pathway targets for controlled mitigation of neuroinflammation long term.

## CONCLUSIONS

In this study, we demonstrated the applicability of Sachi’s Nanoligomer™ platform for high-throughput and rapid neurotherapeutic development for radiation-induced neuroinflammation, through regulation and gene perturbation of a number of upstream immune regulators and canonical pathways. First, we evaluated several Nanoligomer up- and down-regulators using human astrocytes and PBMC. Then, we demonstrated that neuroinflammation caused by radiation exposure in mouse model can be reversed through identification of single key target (GM-CSF) identified here. Using a novel transcriptional inhibitor targeting GM-CSF (CSF2), it was possible to not only treat specific protein expression dysfunction (caused by radiation) but downregulate all key neuroinflammation cytokines in mouse brain tissue. The Nanoligomer induced reversible gene expression modulation can have important therapeutic application by combining specificity and ease of delivery, with the ability to target key gene networks to alleviate dysfunction in a number of key protein expression. This represents a both strong key target ID platform, as well as a therapeutic modality.

## MATERIALS AND METHODS

### Nanoligomer Design and Synthesis

DNA and transcript sequences of gene targets of interest were used as input to Sachi’s bioinformatics toolbox. The gene sequences/transcripts (different numbers based on location, covering different regions of target gene) were interrogated for optimal design based on sequence identity, predicted gene regulatory sites, thermodynamic binding optimization, self-complements, off target effects, solubility, and synthesis parameters. Different TNFR1 and other variants were annotated in the NCBI genome database. Further, machine-learning based on Naïve-Bayes classifiers and experimental data was used to rank, and then validate, the different candidates for gene-perturbation of the potential target.^49–52^ This analysis was used to select the Nanoligomer candidates with the least off-target effects. Following design and ranking, a Nanoligomer was selected and synthesized using our high throughput, automated peptide synthesizer, AAPPTEC Vantage (AAPPTEC, LLC) with solid-phase Fmoc chemistry at a 10-µmol scale on 4-methylbenzhydrylamine (MBHA) rink amide resin. Fmoc–Nanoligomers monomers were obtained from PolyOrg Inc., with A, C, and G monomers protected with Bhoc groups. Purified Nanoligomers’ were quantified by UV-Vis (Thermo Scientific NanoDrop) according to known optical parameters and then stored at −20 °C until further use.

### Astrocyte culture and treatments with Nanoligomers

Primary human astrocytes (Lonza) were maintained in complete astrocyte growth medium (ScienCell) at 37 °C and 5% CO_2_ in a humidified incubator and subcultured at ∼80-90% confluency. Stock solutions of the cytokines (TNF-α, IL-1α, C1q-Sigma-Aldrich) were prepared in molecular biology grade water, aliquoted and stored at −80°C. Composition of cytokine cocktail for astrocyte activation was: TNF-α : 0.03 ng/μL, IL-1α : 0.30 ng/μL and C1q: 4 ng/μL (effective concentrations in the culture medium).^4,57^ For nanoligomer treatment, 10,000 cells were seeded per well of a 48-well plate and cultured until 80% confluency. Then, astrocytes were first pre-treated with the gene-specific Nanoligomers (with or without cytokine cocktail) for 24 hours, after another 24 hours media supernatants were collected and analyzed for secreted protein/cytokine expression.

### PBMC treatment with Nanoligomers

Human PBMCs (Zen Bio, Inc, Research Triangle, NC) were thawed per manufacturer’s instructions and suspended in complete medium (RPMI-1640 w/ Glutamax + 10% heat inactivated fetal bovine serum (FBS) + 1% Penicillin /Streptomycin. PBMCs (500k/190 µL per well of a U-bottom 96-well plate) were then treated with 10 µL (stock conc: 200µM in Nanoligomer media) of the nanoligomer solution and cultured for 24 hours at 37 °C and 5% CO2 in a humidified incubator after which they were exposed to various radiation doses (3 Gy, Control: 0 Gy, no radiation). Nanoligomer media (vehicle negative control) was added instead of Nanoligomer for no treatment control. After 24 hours, the cell culture supernatants were collected and stored at − 80°C, and cells were immediately stained for flow cytometry analysis.

### Quantification of secreted proteins using Immune Monitoring 65-Plex Human ProcartaPlex™ Panel

Cell culture supernatants were analyzed for secreted proteins using Immune Monitoring 65-Plex Human ProcartaPlex™ Panel for MAGPIX (Thermofisher Scientific, Carlsbad, CA) as per manufacturer’s instructions. This is a preconfigured multiplex immunoassay kit that measures 65 cytokines, chemokines, and growth factors for efficient immune response profiling, biomarker discovery, and validation. It consists of two separate kits: a 43-plex and a 22-plex kit that have been specifically designed to be used on the MAGPIX instrument that can detect up to 50 targets simultaneously. In this study, PBMC supernatants were probed for the following 65 markers: APRIL, BAFF, BLC, CD30, CD40L, ENA-78, Eotaxin, Eotaxin-2, Eotaxin-3, FGF-2, Fractalkine, G-CSF, GM-CSF, GROα, HGF, IFN-α, IFN-ɣ, IL-1α, IL-1β, IL-2, IL-2R, IL-3, IL-4, IL-5, IL-6, IL-7, IL-8, IL-9, IL-10, IL-12p70, IL-13, IL-15, IL-16, IL-17A, IL-18, IL-20, IL-21, IL-22, IL-23, IL-27, IL-31, IP-10, I-TAC, LIF, MCP-1, MCP-2, MCP-3, M-CSF, MDC, MIF, MIG, MIP-1α, MIP-1β, MIP-3α, MMP-1, NGF-β, SCF, SDF-1α, TNF-α, TNF-β, TNF-R2, TRAIL, TSLP, TWEAK, and VEGF-A. The plate was read using the MAGPIX instrument and xPONENT® software (version 4.2, Luminex Corp, Austin, Texas, US). The overall intra- and inter-assay precision is reported by the manufacturer as 2–19%, and accuracy as 87–107% over the calibration range of 3.2–10,000 pg/mL protein concentration. Data analysis was performed in Microsoft Excel and plots were made using OriginLab software.

### RNA extraction and real-time quantitative polymerase chain reaction (RT-qPCR)

RNA was extracted using a miRNeasy Micro Kit (QIAgen, USA) per manufacturer protocol and reverse transcribed to cDNA using High-Capacity cDNA Reverse Transcription Kit (Applied Biosystems). qPCR was performed using Fast SYBR Green Master Mix (Applied Biosystems) and custom intron-spanning primers (Table S1, purchased from Integrated DNA Technologies, Inc.) on Bio-Rad CFX96 Touch Real-Time PCR Detection System. Gene expression of *CSF2* was analyzed following the delta-delta Ct method relative to the reference gene, Glucuronidase Beta, *GUSB*.

**Table S1.**
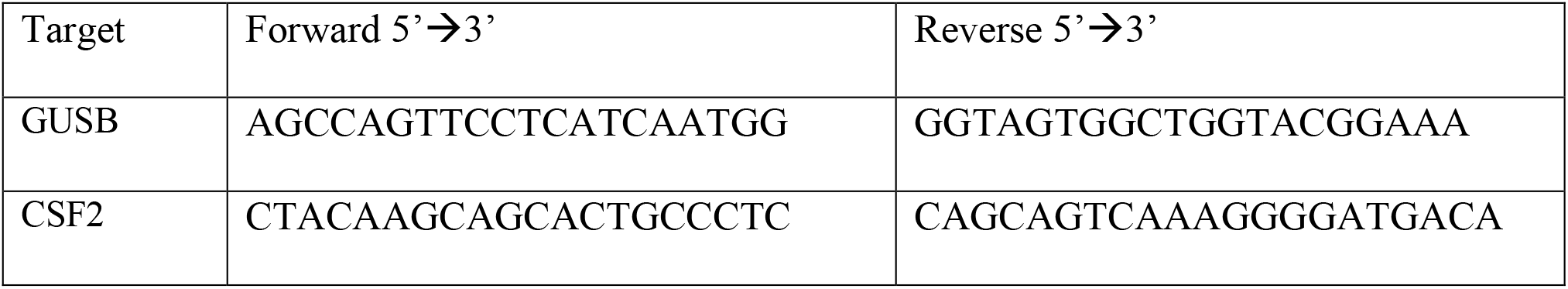
Primers used for human qPCR.

### Murine model study

The animal care facilities at Colorado State University (CSU) are AAALAC accredited, and all animal work was approved by the CSU Institutional Animal Care and Use Committee under protocol 1891. Female C57BL/6NCrl mice (Charles River Laboratories) were purchased at 6 weeks of age and acclimated for 2 weeks prior to experimental start. Three groups received intraperitoneal (IP) injections of PBS (Groups 1 and 2) or 201.31U.1_CSF2 Nanoligomer at 3 mg/kg (Group 3) (see supplemental Fig. S12). 24 hours post-injection, the mice were irradiated to the whole body with 3 Gy of ^137^Cs γ-rays at a dose rate of 0.78 Gy/minute (Fig. 4A). Four hours later, the mice were euthanized by CO_2_ inhalation and blood was collected by cardiac puncture. Whole blood was processed to serum which was frozen. Necropsy continued with collection of whole brain which was then dissected under a microscope to collect only cortex. All samples were immediately flash frozen and stored at −80°C.

### Quantification of Cytokine & Chemokine in mouse tissues

Flash frozen tissues were homogenized using a mortar and pestle and syringe disruption to form a homogenate solution in tissue cell lysis buffer (EPX-99999-000). Samples were then centrifuged at 16,000xg for 10 min at 4°C and supernatant was transferred to a fresh tube. Bio-Rad’s DC Protein Assay Kit was used to determine protein content and all samples were diluted to 10 mg protein/mL. Quantification of cytokines/ chemokine was performed with Cytokine & Chemokine Convenience 36-Plex Mouse ProcartaPlex Panel 1A (EPXR360-26092-901), analyzed on a Luminex MAGPIX xMAP instrument, and quantified using xPONENT software. For standard curves, eight four-fold dilutions of protein standards were used. Target list includes: ENA-78 (CXCL5), Eotaxin (CCL11), GRO alpha (CXCL1), IP-10 (CXCL10), MCP-1 (CCL2), MIP-1 alpha (CCL3), MIP-1 beta (CCL4), MIP-2 alpha (CXCL2), RANTES (CCL5), G-CSF (CSF-3), GM-CSF, IFN alpha, IFN gamma, IL-1 alpha, IL-1 beta, IL-2, IL-3, IL-4, IL-5, IL-6, IL-9, IL-10, IL-12p70, IL-13, IL-15/IL-15R, IL-17A (CTLA-8), IL-18, IL-22, IL-23, IL-27, IL-28, IL-31, LIF, MCP-3 (CCL7), M-CSF, TNF alpha.

## ACKNOWLEDGMENTS

This work was supported by financial support from National Aeronautics and Space Administration SBIR Contracts 80NSSC21C0242 and 80NSSC22CA116 to Sachi Bioworks, and funding from NASA grant NNX15AK13G to MMW. The authors thank Prof. Alexander Brandl, Director, Irradiation Services Laboratory and Justin Bell at Colorado State University for their time spent in coordination and irradiation of PBMC samples, and Colleen Courtney for helping with mouse tissue ProcartaPlex sample preparation.

---

**Table 1:** Prioritized gene targets for neuroinflammation countermeasure development. Italics of Approved/In Trial Drug indicates that the mechanism of action of the drug on target gene is proposed to be the opposite needed for immune dysregulation correction

| Target Gene | Target Function | *Approved/In Trial Drug | Ref. |
| --- | --- | --- | --- |
| Granulocyte-Macrophage Colony-Stimulating Factor (CSF2) | Cytokine, Growth factor that controls the production, differentiation, and function of granulocytes and macrophages | *Sargramostim/Leukine\**<br>*Molgramostim* | 18,75–78 |
| Interleukin-6 (IL-6) | Cytokine with a wide variety of biological functions in immunity, tissue regeneration, and metabolism. | *Tocilizumab* | 18,76–79 |
| IL-10 | Prominent role in limiting immune-mediated inflammation during radiation exposure, infection, allergy, or autoimmunity |  | 46 |
| Interleukin-1β (IL-1β) | Potent proinflammatory cytokine. Initially discovered as the major endogenous pyrogen, induces prostaglandin synthesis, neutrophil influx and activation, T-cell activation and cytokine production, B-cell activation and antibody production, and fibroblast proliferation and collagen production. | *Anakinra* | 18,76–78,80–83 |
| Tumor Necrosis Factor-α (TNF-α) | Cytokine that binds to TNFRSF1A/TNFR1 and TNFRSF1B/TNFR2. Mainly secreted by macrophages and can induce cell death of certain tumor cell lines. | *Infliximab (Remicade) Adalimumab (Humira) Certolizumab pegol (Cimzia)/ Etanercept (Enbrel)* | 18,76–78,84 |
| TNFR1 | Overexpression promotes apoptosis and activates NF-κB, which leads to proinflammatory cytokine production and glial activation, leads to neuroinflammation and neuronal death | | 4,43 |

