## Supplementary Information for "Screening Immunotherapy Targets to Counter Radiation-Induced Neuroinflammation"

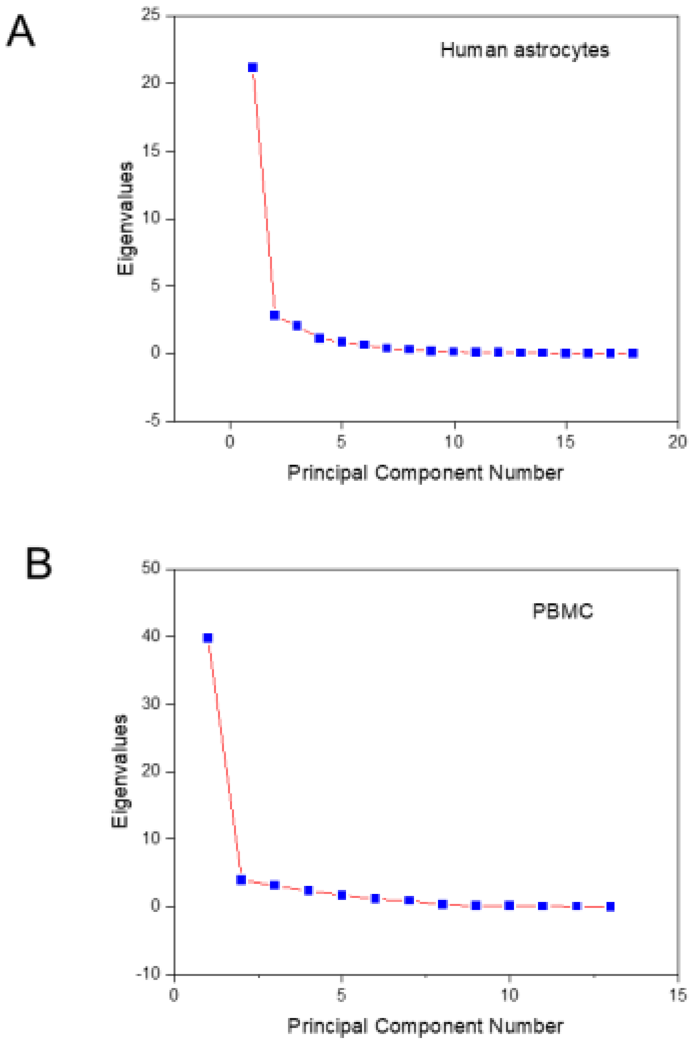


**Figure S1.** Scree plots of principal component analysis for (A) human astrocytes and (B) peripheral blood mononuclear cell (PBMC) screens using multiplex cytokine panel. These plots indicate that PC1 captures most of the variability in the data for both cell models.

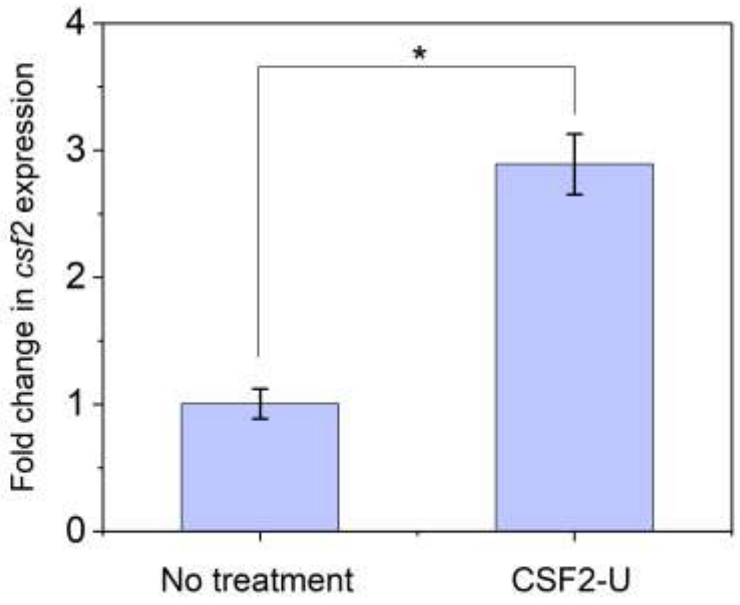


**Figure S2.** Nanoligomers regulate expression of GM-CSF in HepG2 Cells. (A) mRNA expression of GM-CSF (csf2) is upregulated at the transcriptional level when HepG2 cells are treated with CSF2-U. Data shown represents five biological replicates (* $p<0.001$).

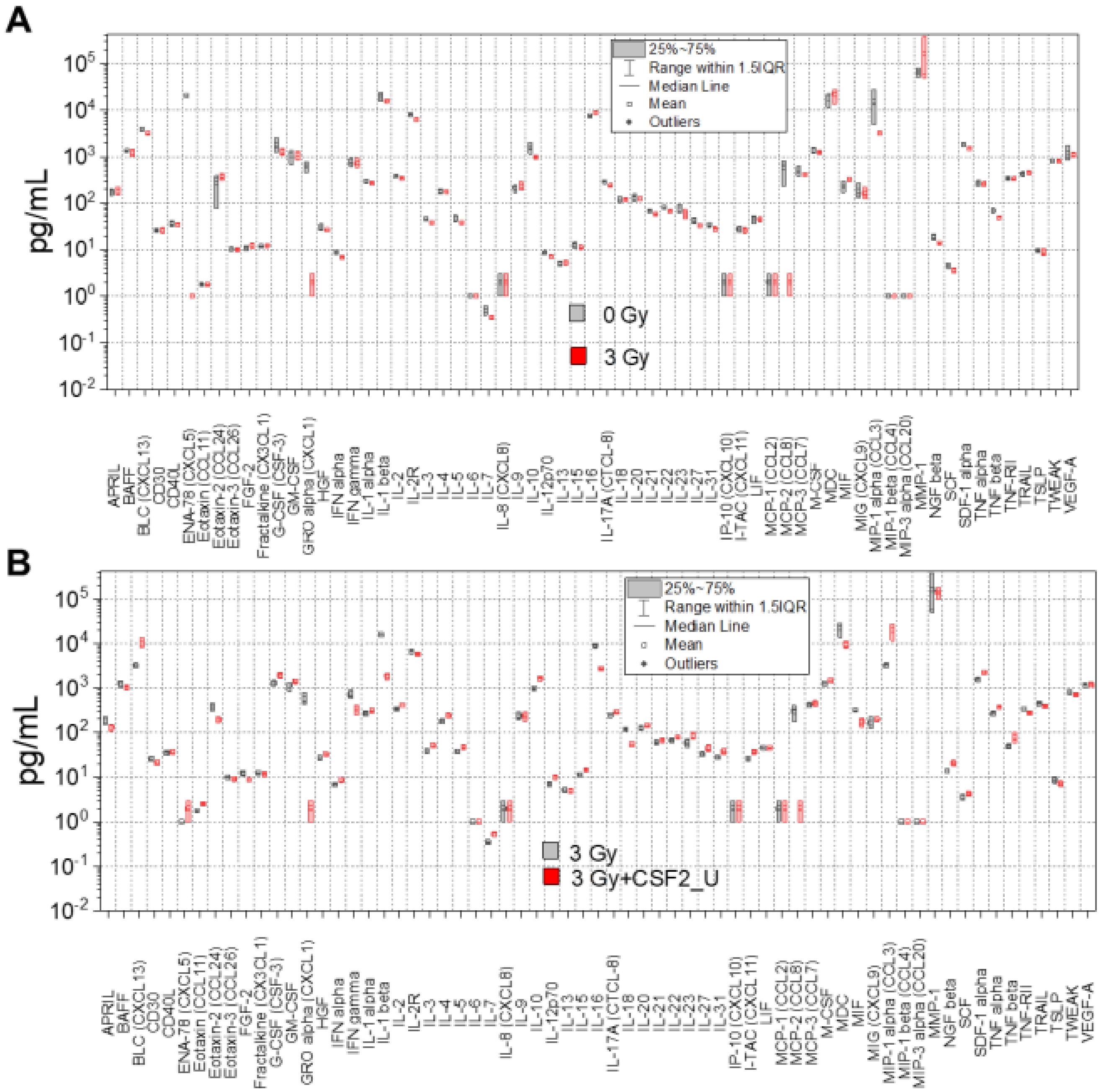


**Figure S3.** Nanoligomers regulate cytokine expression. (A) Cytokine expression in absence (0 Gy) and presence (3 Gy) of Gamma radiation. (B) Cytokine expression in presence of Gamma radiation (3 Gy) without and with Nanoligomer treatment of GM-CSF upregulator (CSF2_U) and. Data shown represents three biological replicates.

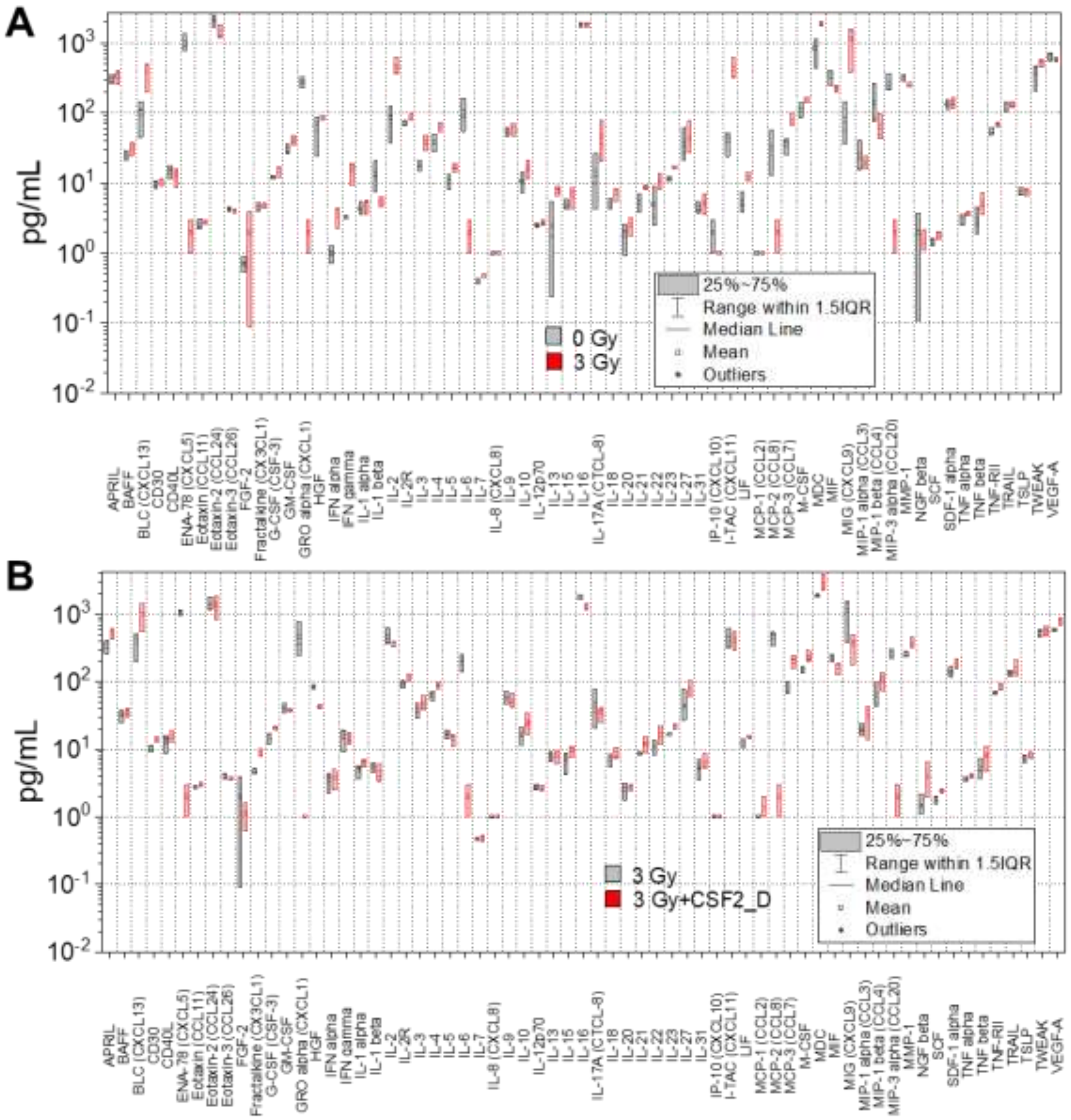


**Figure S4.** Nanoligomers regulate cytokine expression. (A) Cytokine expression in absence (0 Gy) and presence (3 Gy) of Gamma radiation. (B) Cytokine expression in presence of Gamma radiation (3 Gy) without and with Nanoligomer treatment of GM-CSF downregulator (CSF2_D) and. Data shown represents three biological replicates.

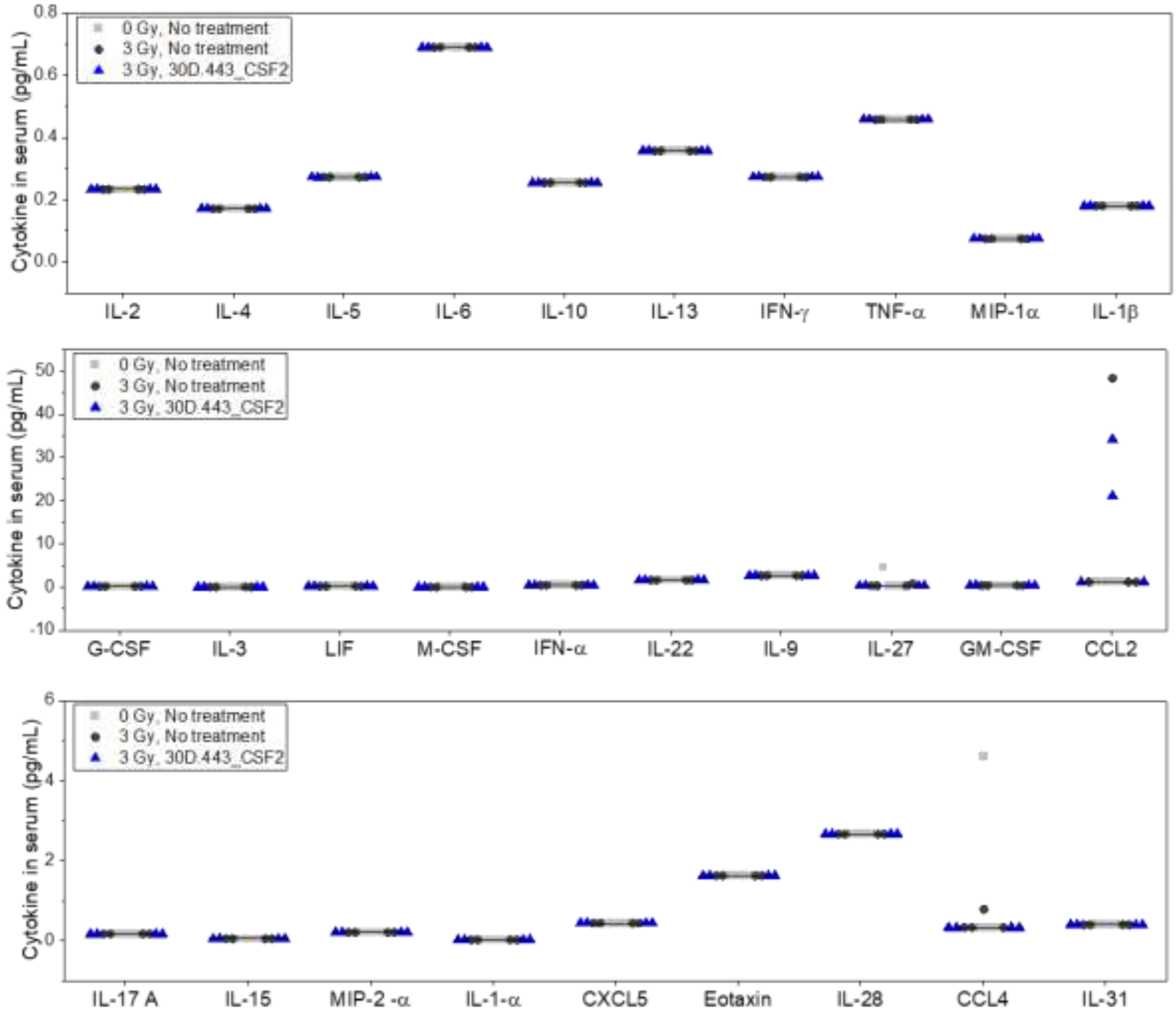


**Figure S5.** Sachi's Nanoligomer 30D.443_CSF2 is safe and non-immunogenic *in vivo*. Commonly modulated cytokines associated with pathological responses including acute-phase response, cytokine storm/release, fibrosis, hemophagocytic syndrome, neutrophilic inflammation, systemic inflammatory response syndrome, and Th1 and Th2 immune response showed no significant increase in response to 30D.443_CSF2 treatment. Mostly, the cytokines were below the limit of detection. CCL2 was elevated in two mice with Nanoligomer treatment (3 Gy) and one mouse with vehicle (3 Gy), which could be due to radiation exposure.

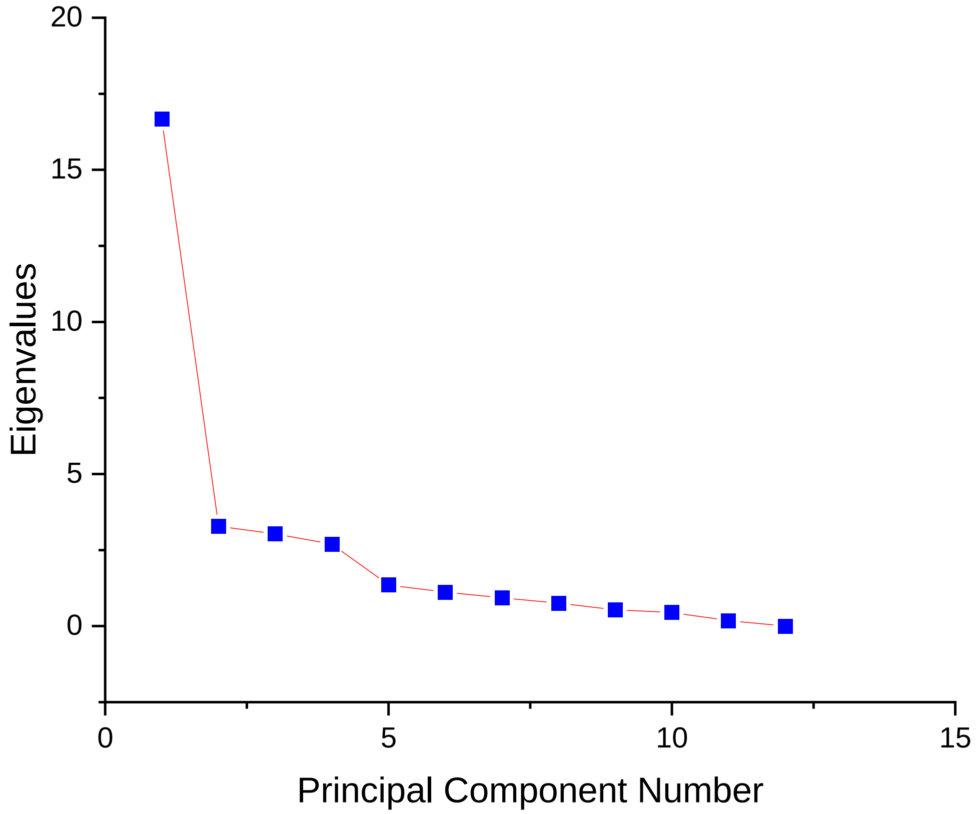


**Figure S6.** Scree plot of the principal component analysis of the mouse cytokine data (0 Gy, 3Gy radiation, 30D.443_CSF2 nanoligomer) using multiplex cytokine panel. Principal component analysis of the cytokine data revealed that principal component 1 (PC1) capture the most of the variability in the data. PC2, PC3, and PC4 capture only 10.59%, 9.79%, and 8.69% of variability, respectively.

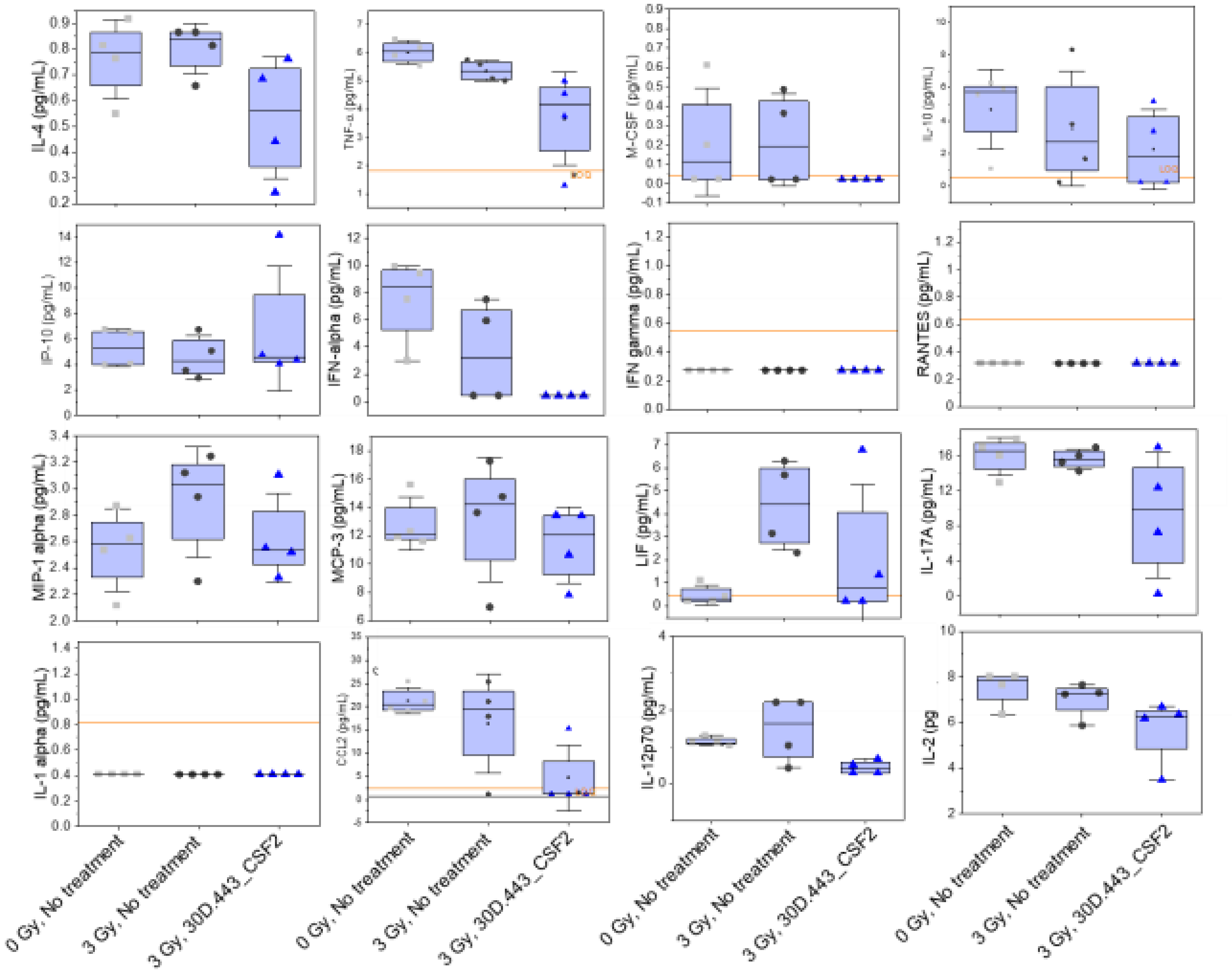


**Figure S7.** Cytokine level in the cortex of mice with and without 3 Gy radiation and treatment with 30U.443_CSF2. Orange line represented the limit of quantification of the assay, symbols represent individual animals, horizontal bar is the average and error bars are +/- one standard deviation.

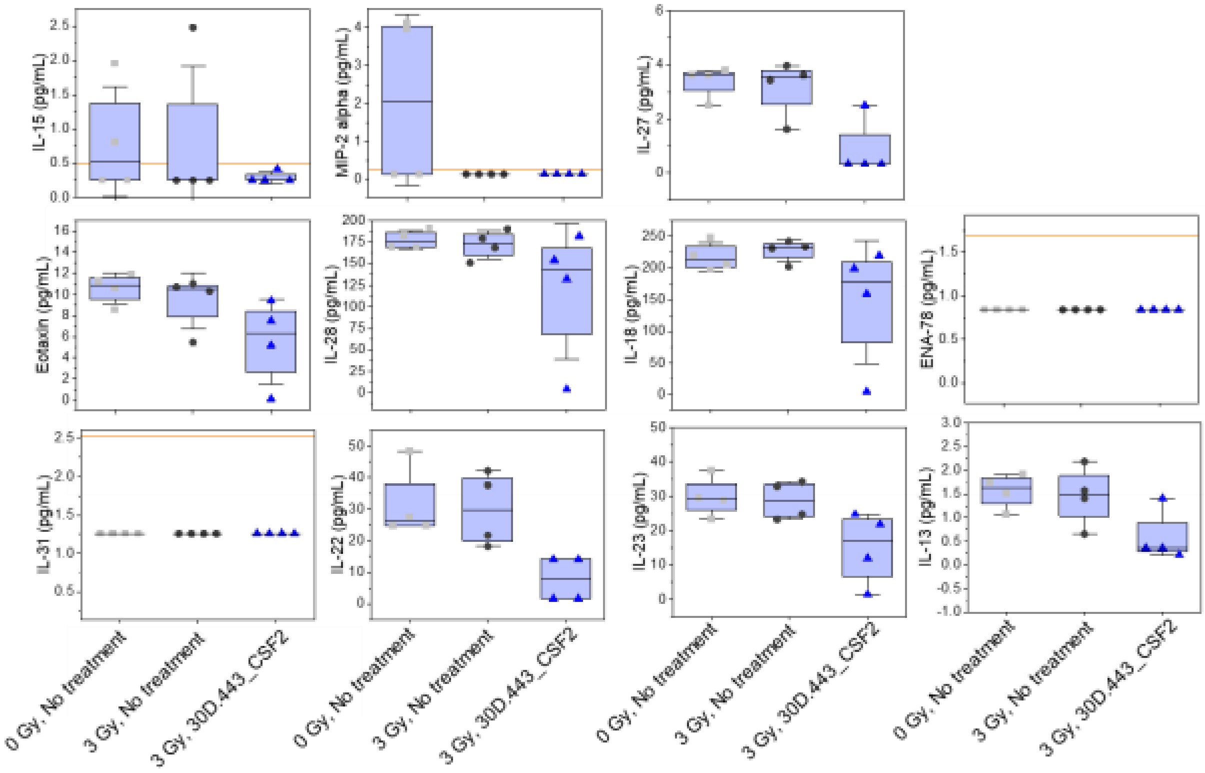


**Figure S7 Continued.** Cytokine level in the cortex of mice with and without 3 Gy radiation and treatment with 30U.443_CSF2. Orange line represented the limit of quantification of the assay, symbols represent individual animals, horizontal bar is the average and error bars are +/- one standard deviation.
